# Identification and characterization of functional and trans-dominant negative HERV-K (HML-2) Rec proteins encoded in the human genome

**DOI:** 10.1101/2025.07.07.663474

**Authors:** Katarzyna Zurowska, Godfrey Dzhivhuho, David Grabski, David Rekosh, Marie-Louise Hammarskjold

**Affiliations:** Myles H. Thaler Center for AIDS and Human Retrovirus Research, School of Medicine, University of Virginia, Charlottesville, Virginia, USA; Department of Microbiology, Immunology, and Cancer Biology, School of Medicine, University of Virginia, Charlottesville, Virginia, USA; Department of Surgery, School of Medicine, University of Virginia, Charlottesville, Virginia, USA

## Abstract

Human Endogenous Retroviruses K (HERV-K) of the HML-2 subgroup are the most recently integrated and biologically active retroviral elements within the human genome. The HERV-K Rec protein, a functional homolog of HIV Rev and HTLV Rex, is necessary for the nuclear export of viral mRNAs with retained introns. However, the diversity of Rec proteins encoded in the human genome and their functional capacities have remained largely unexplored. We identified full-length Rec protein sequences in the human genome and selected 23 variants for functional characterization. Using a dual-color fluorescent reporter and a complementary ELISA assay, we found that Rec proteins from only 7 genomic loci were functional. In addition, a subset of the non-functional Rec proteins exerted trans-dominant negative effects. Detailed mutational analysis of the most potent inhibitory variant, encoded by the HERV-K provirus 12q14.1, displayed only 2 amino acid changes (H2N and del34E) relative to the prototypical functional Rec protein encoded by the reconstructed consensus HERV-K (HERV-K Con). Insertion of a glutamic acid at position 34 restored the functional activity, while the substitution of histidine with asparagine at position 2 did not. Our results unveil an unexpected complexity in HERV-K (HML-2) post-transcriptional regulation, with Rec variants displaying a spectrum of activities ranging from robust nuclear export of RNA with retained introns to trans-dominant negative inhibition of Rec function. These findings expand our understanding of the regulatory landscape governing HERV-K (HML-2) expression and suggest mechanisms by which different HERV-K Rec proteins may influence host cell biology and pathology, including oncogenesis.

## Introduction

Human Endogenous Retroviruses (HERVs), which constitute about 8% of the human genome, are a class of transposable elements that are remnants of ancient exogenous retroviruses. These viruses integrated into the human germline between 100 and 40 million years ago (1, 2). Although HERVs have accumulated many mutations and deletions within coding sequences over time (3), many of these elements remain transcriptionally active, contributing to cellular function either through the production of viral proteins or by serving as regulatory elements for host gene expression (3–6).

HERV taxonomy is based on sequence similarity to other animal retroviruses (7–9). The HERV-K clade belongs to class II of beta retrovirus-like endogenous retroviruses, with the "K" designation referring to the lysine tRNA primer used to prime reverse transcription in these viruses. The HERV-K (HML-2) subgroup that comprises more than 90 proviruses is the most recently integrated and biologically active. In this report, HERV-K (HML-2) virus will hereafter be referred to simply as HERV-K.

Many HERV-K copies retain intact open reading frames (ORFs) with the potential to produce viral RNAs and, in many cases viral proteins. Complete HERV-K genomes, like all intact retroviruses, encode the essential gag, pro, pol, and env genes, flanked by long terminal repeats (LTRs) (7). Additionally, like the more complex viruses, such as HTLV-1 and HIV, they also encode regulatory proteins (10). HERV-K proviruses are classified as type 1 or type 2. Type 2 proviruses encode the Rec protein from a doubly spliced transcript; type 1 proviruses have mutations and a 292bp deletion in the pol-env region, that deletes the first coding exon of *rec* and changes the doubly spliced transcript in some of the type 1 viruses to encode a protein named Np9 (7, 10–12).

Rec interacts directly with the Rec-Response Element (RcRE), present in the 3’ end of all HERV-K mRNAs (10, 13, 14). Through this interaction, Rec facilitates the nucleocytoplasmic export of HERV-K mRNAs that contain retained introns, using the Crm1-RanGTP pathway (10, 15). Rec contains an arginine-rich nuclear location signal (NLS), as well as a nuclear export signal (NES) (15, 16), and functions similarly to the HIV Rev, HTLV Rex, and MMTV Rem proteins (17–22). In addition to the proviruses, more than 900 HERV-K solitary LTRs are scattered throughout the human genome (5) . Many of these contain RcRE sequences that can potentially interact with Rec proteins. Recent studies have identified cellular genes that contain sequences with high homology to HERV-K RcREs, suggesting that Rec may modulate cellular gene expression through interactions with these elements (5, 23–26).

Some evidence suggests that HERV-K Rec may play significant roles in cancer development, potentially through interaction with cellular proteins. For example, Rec has been shown to interact with the promyelocytic leukemia zinc-finger protein (PLZF), activating c-myc proto-oncogene expression and promoting cell proliferation (12, 27). Rec has also been shown to interact with the human small glutamine-rich tetratricopeptide repeat-containing protein (hSGT). This leads to increased androgen receptor activity, which may promote oncogenesis (28).

The first coding exon of Rec is the same as the first 87 amino acids of the 95 amino acid Env signal peptide (SP) (29). The second coding exon also overlaps *env* sequences but is translated in a different open reading frame. In the case of the closely related Mouse Mammary Tumor Virus (MMTV), it has been shown that the Env SP, which corresponds to the first coding exon of the nuclear export protein Rem, is sufficient to function in the nuclear export of viral mRNAs (30) since it contains RNA binding, as well as nuclear import (NLS) and export (NES) domains. The HERV-K Env SP also contains the NLS and NES sequences, which suggests that it could also be sufficient for HERV-K mRNA export. This relationship and the fact that both coding exons of Rec overlap the *env* coding region become particularly relevant when interpreting studies showing oncogenic properties of HERV-K Env in various cancers (31, 32), as Env expression is also likely to depend on Rec expression.

Despite the widespread distribution of HERV-K sequences in the human genome and the potential importance of Rec in regulating viral and cellular gene expression, an analysis of which proviral loci produce Rec proteins that can function in RNA export has not been conducted. In this study, we performed a comprehensive functional analysis of HERV-K Rec proteins from many different proviral loci. Our findings reveal that while some Rec variants retain robust nuclear export of mRNA with retained introns, others act as trans-dominant negative regulators, suggesting that Rec regulation of HERV-K expression is complex.

## Results

### Characterization and Selection of HERV-K HML-2 Rec Proteins for Functional Analysis

The identification and characterization of functional HERV-K HML-2 Rec proteins present significant challenges due to the repetitive nature and abundance of HERV sequences in the human genome. We approached this problem by first annotating key proviral features of the reconstituted HERV-K provirus, HERV-K Con, including long terminal repeats (LTRs), open reading frames (ORFs), and known splice sites (Figure 1A) (33, 34). To account for both type 1 and type 2 proviruses, we generated two reference genomes: the original HERV-K Con (type 2) and a modified version with a 292bp deletion in the pol-env region (type 1). We then aligned 91 HERV-K genomes (5) to these annotated references, transferring annotations to each sequence. This process enabled us to differentiate between type 2 proviruses (56 copies) containing the *rec* gene and type 1 proviruses with the 292bp deletion. We then extracted and joined the *rec* exons from the type 2 viruses according to the known splicing pattern, translated the 56 Rec coding sequences, and aligned them to the HERV-K Con Rec protein (Figure 1B).

**Figure 1:**
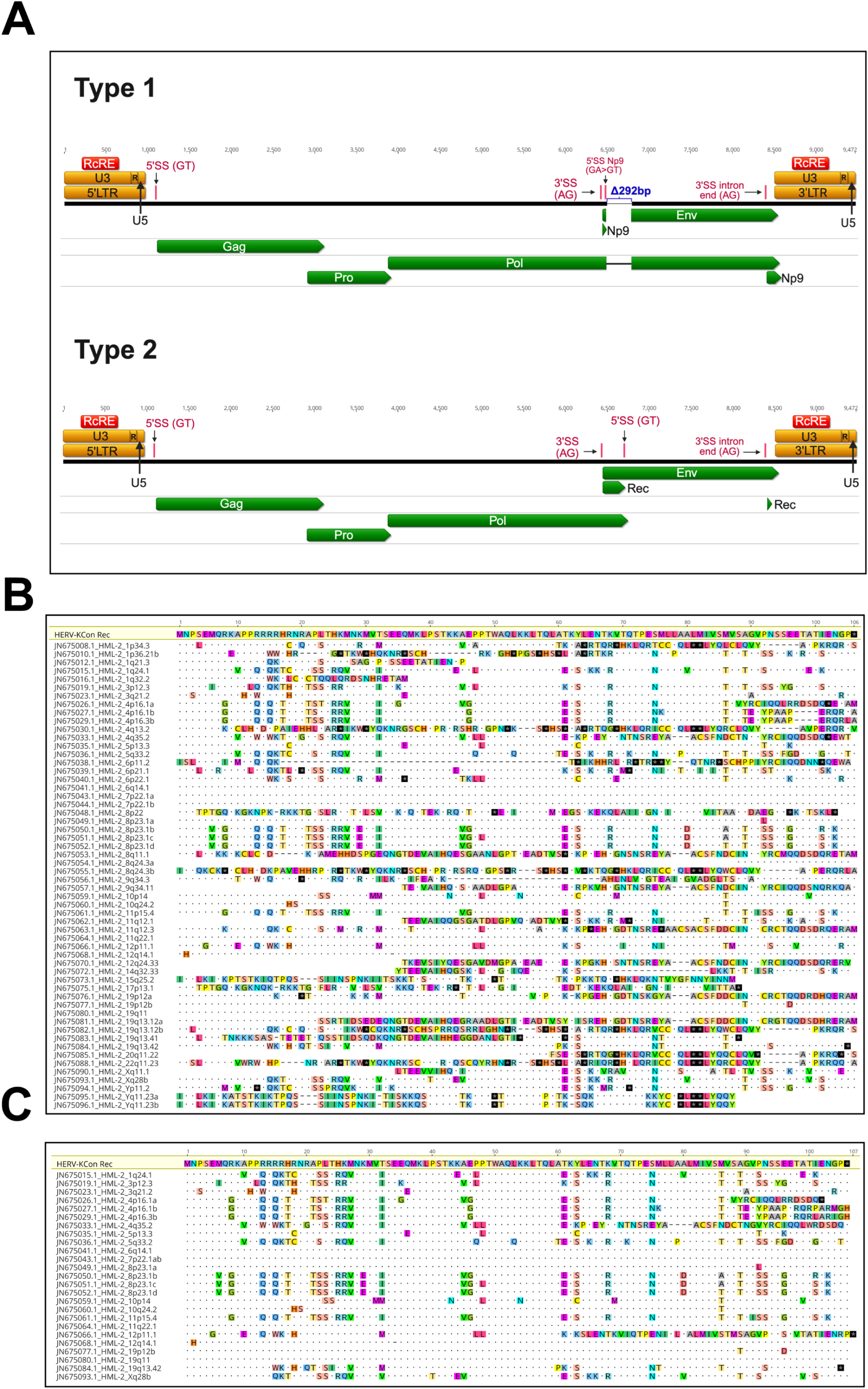
Bioinformatic characterization and selection of HERV-K HML-2 Rec Proteins for functional analysis. (A) Diagram of the primary features of the HERV-K Con provirus, including long terminal repeats (LTRs), open reading frames (ORFs), and specific splice sites. This schema was used for the annotation of type 1 (modified version with a 292 bp deletion in the pol-env junction: top panel) and type 2 (the original HERV-K Con: bottom panel) proviral reference genomes. Relative to type 2, there is an A to T point mutation in many type 1 viruses that introduces a stop codon in the env reading frame. This also creates a new 5’ splice site, which allows for the generation of a doubly spliced mRNA that encodes the Np9 protein. Eight bases downstream from the splice site, the 292 bp deletion removes most of the first coding exon of rec. A downstream methionine in type 1 viruses may allow for translation of a N-terminally truncated Env protein. This protein would not be expected to be glycosylated, as it lacks a signal peptide. (B) Alignment of translated Rec coding sequences (extracted and joined using the known splice sites in type 2 proviruses) to the HERV-K Con Rec protein. This process yielded 56 distinct Rec protein sequences. Differences relative to the HERV-K Con Rec protein sequence are highlighted in color. Dots represent identical amino acid residues, while dashes represent a deletion. Stop codons are highlighted as black squares with a * symbol. (C) Comparative sequence analysis of the 26 full-length Rec proteins. Differences relative to the HERV-K Con Rec sequence are highlighted in color. Dots represent identical amino acid residues, while dashes represent a deletion. Stop codons are highlighted as black squares with a * symbol.

This alignment revealed striking protein sequence diversity among the Rec proteins encoded by these HERV-K loci (Figure 1B). Over half of the ORFs were truncated due to large deletions and/or premature stop codons. We identified and selected 26 full-length Rec proteins, excluding the 30 incomplete or heavily truncated protein sequences from further analysis. Among these 26 full-length proteins, we observed significant sequence diversity compared to the reference HERV-K Con Rec (Figure 1C). However, four Rec proteins (encoded by proviruses 6q14.1, 7p22.1ab, 11q22.1, and 19q11) were identical to the reference HERV-K Con Rec at the protein level, despite 1-3 nucleotide differences in their coding sequences (34). Thus, we selected 23 distinct Rec protein variants for functional analysis: one identical to the reference HERV-K Con Rec and 22 with amino acid variations.

### Evaluation of HERV-K HML-2 Rec Protein Activity using an RcRE Reporter Assay

To evaluate the functional capacity of the selected HERV-K HML-2 Rec proteins, we developed an assay using a dual-color fluorescent reporter vector. This vector is similar to our established HIV-1 Rev-RRE dual-color reporter, but contains a tandem RcRE in place of the HIV RRE (35, 36). This modified reporter produces mCherry constitutively from a spliced mRNA, while GFP expression from an unspliced RNA depends on the presence of both the cis-acting RcRE and a functional trans-acting Rec protein. To facilitate our experiments, we generated a 293T/17-based cell line stably expressing the reporter mRNA (Figure 2A).

**Figure 2:**
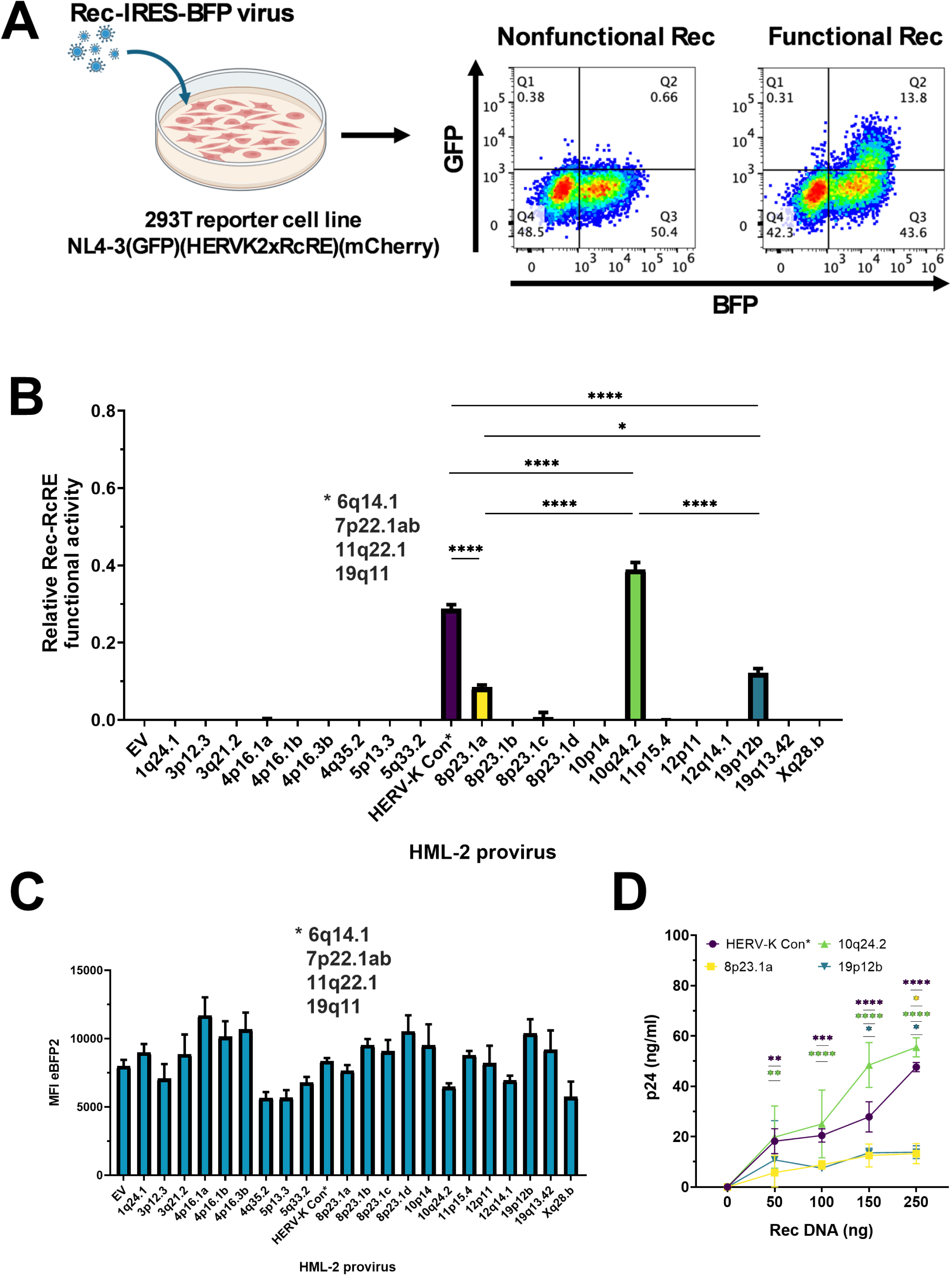
Functional Analysis of HERV-K Rec Proteins. (A) Schematic of the dual-color Rec-RcRE reporter assay using retroviral vectors expressing Rec and eBFP2. The cells contain a reporter vector that constitutively produces mCherry from a fully spliced transcript. Upon transduction with the Rec-expressing vectors, the mCherry-positive cells also express eBFP2. Functional Rec proteins interact with the RcRE, promoting nuclear export of the unspliced mRNA encoding GFP (right flow chart). Non-functional Rec proteins result in only eBFP2 expression (left flow chart). (B) Quantification of Rec functional activity using the dual-color reporter assay by flow cytometry analysis performed 72 hours post-transduction. 293T RcRE reporter cells were transduced with vectors expressing the various Rec proteins and eBFP2. Rec activity was assessed by gating on mCherry-positive cells, selecting eBFP2-positive cells (indicating Rec vector expression), and measuring GFP and eBFP2 expression. Rec-RcRE activity was calculated as the ratio of GFP mean fluorescence intensity to eBFP2 mean fluorescence intensity. The HERV-K Con Rec sequence represents four identical Rec proteins (6q14.1, 7p22.1ab, 11q22.1, 19q11) (marked with an asterisk). The data is shown as the mean value ± SD from three independent experiments. Significant differences were assessed by ordinary One-Way ANOVA with Tukey’s post-hoc test (*p<0.05, ****p<0.0001). (C) Transduction of the 293T/17 RcRE reporter cells with Rec and eBFP2 expressing vectors. The MFI of eBFP2 expressed from each of the individual transductions in (B) was determined using flow cytometry. Data shown as mean ± SD from three independent experiments. (D) Validation of Rec activity using a p24 release assay. 293T/17 cells were co-transfected with equal amounts of a Gag-Pol-RcRE reporter vector (1500 ng) and increasing amounts of functional Rec vectors (50, 100, 150, 250 ng). An empty vector was used to normalize the total DNA input to 2000 ng. Supernatants were collected at 72 hours post-transfection and p24 was then measured by ELISA. The data shown are the mean values ± SD from three independent experiments. Statistical significance was assessed by ordinary Two-Way ANOVA with Dunnett’s multiple comparisons post-hoc test (*p<0.05, **p<0.01, ***p<0.001, ****p<0.0001).

To test the activity of the various Rec proteins, we cloned the ORFs for each of the 23 variants into a murine stem cell virus-derived retroviral vector (pMSCV), positioning them upstream of an IRES-eBFP2 cassette. This bicistronic design ensures simultaneous expression of both Rec and eBFP2 from a single transcript, facilitating detection and quantification of expression levels.

If a functional Rec protein is expressed from this vector in the transduced reporter cells, its interaction with the RcRE results in export of the unspliced mRNA and GFP expression. After flow cytometry analysis, Rec functional activity can be quantified as the ratio of GFP mean fluorescence intensity (MFI) to eBFP2 MFI. By using this ratio, the results are normalized to account for potential differences in transduction efficiency and RNA expression, since Rec and eBFP2 are translated from the same mRNA.

Transduction experiments with the vectors expressing the 23 different Rec proteins showed that in addition to HERV-K Con Rec, encoded by the proviruses at 6q14.1, 7p22.1ab, 11q22.1, and 19q11, only 3 other Rec proteins (at loci 8p23.1a, 10p24.2, and 19p12b) displayed functional activity (Figure 2B). The remaining 19 Rec variants showed no detectable activity, as indicated by the absence of GFP expression despite eBFP2 expression (Figure 2C). We also employed a complementary approach, using an HIV GagPol-RcRE reporter vector to further validate and quantify the activity of the four functional Rec protein variants. This reporter measures p24 release as an indicator of viral protein expression mediated by a functional Rec-RcRE interaction. To perform this assay, the GagPol-RcRE reporter was co-transfected into cells with increasing amounts of the plasmids expressing the Rec proteins (Figure 2D). This confirmed the functionality of all four proteins. Importantly, the rank order of activity was consistent between both assays, suggesting a spectrum of Rec protein functionality, with some variants being more active than others.

We next decided to directly investigate how functional activity was related to steady state levels of Rec expression. Due to the lack of commercially available Rec antibodies, we generated constructs with HA tags added to the N-termini of these Rec proteins, enabling visualization of their expression by Western blot using an anti-HA antibody. As shown in Figure 3A, all four of the tagged constructs displayed Rec activity. Western blot analysis showed some differences in protein expression levels among the HA-tagged Rec variants (Figures 3B and 3C). However, when the Rec activity was normalized based on the protein expression levels (Figure 3D), the rank order of activity was the same as seen with the non-tagged constructs, which had been normalized to eBFP2 expression in Figure 2B.

**Figure 3:**
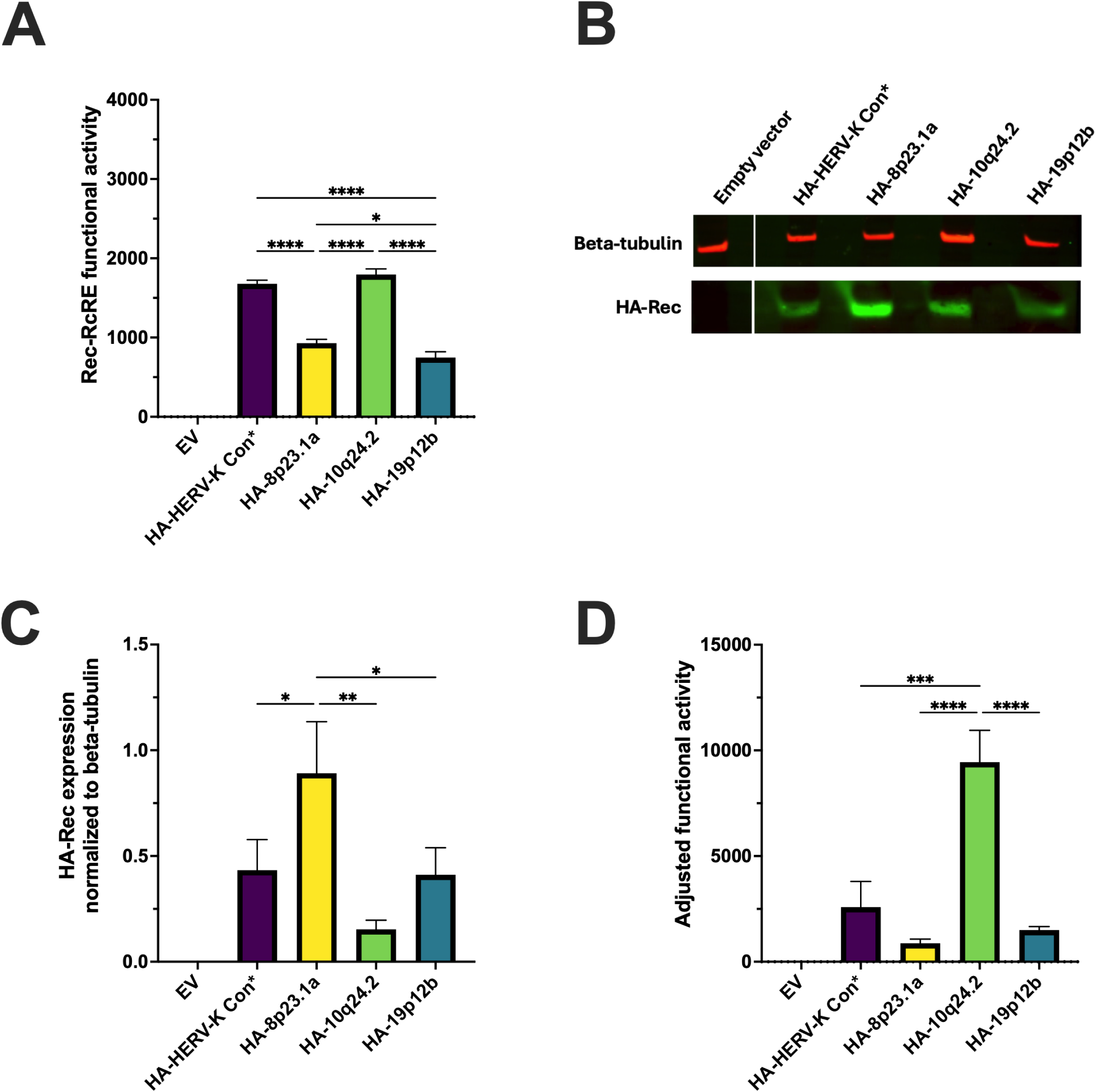
Functional Analysis and Quantitation of HA-Tagged Rec Proteins. (A) Functional activity of HA-tagged Rec variants. 293T RcRE reporter cells were transfected using lipofectamine 3000 with vectors expressing the various Rec proteins. GFP MFI was measured 72 hours post-transfection as an indication of Rec functional activity. HERV-K Con Rec (marked with an asterisk) represents four identical Rec protein sequences (6q14.1, 7p22.1ab, 11q22.1, 19q11). Statistical significance assessed by ordinary One-Way ANOVA with Tukey’s post-hoc test (*p<0.05, **p<0.01, ***p<0.001, ****p<0.0001). (B) Western blot analysis of HA-tagged Rec protein expression. Upper panel: β-tubulin loading control; lower panel: HA-tagged Rec detection. Representative Western blot from two independent experiments using two different plasmid preparations. (C) Quantification of HA-tagged Rec protein expression levels normalized to β-tubulin loading control. Data represent the mean value ± SD from two independent experiments with triplicate samples using two different plasmid preparations. Statistical significance assessed by ordinary One-Way ANOVA with Tukey’s post-hoc test (*p<0.05, ***p<0.001). (D) Functional activity of HA-Rec variants normalized to protein expression. Rec-RcRE functional activity (from panel A) was normalized to HA-tagged protein expression levels (panel C) by calculating the ratio of GFP MFI to normalized HA-Rec levels. Data represents the mean value ± SD from 2 independent experiments with triplicate samples using two different plasmid preparations.

### Some HERV-K HML-2 Rec Proteins are trans-dominant negative

We hypothesized that some of the non-functional Rec proteins might display a trans-dominant negative phenotype. To test this, we selected the four non-functional Rec proteins (encoded at 3q21.2, 5p13.3, 10p14, and 12q14.1) that were the least changed (< 15 aa changes) compared to HERV-K Con Rec (Figure 4A). We cloned cDNA copies of each coding sequence, with or without N-terminal HA-tags, into expression plasmids. The HA-tagged plasmids were then transfected into the RcRE reporter cell line. Western blot analysis revealed that all four selected non-functional Rec proteins were expressed in these cells, but the Rec protein encoded by provirus 3q21.2 showed only very low levels (Figure 4B).

**Figure 4:**
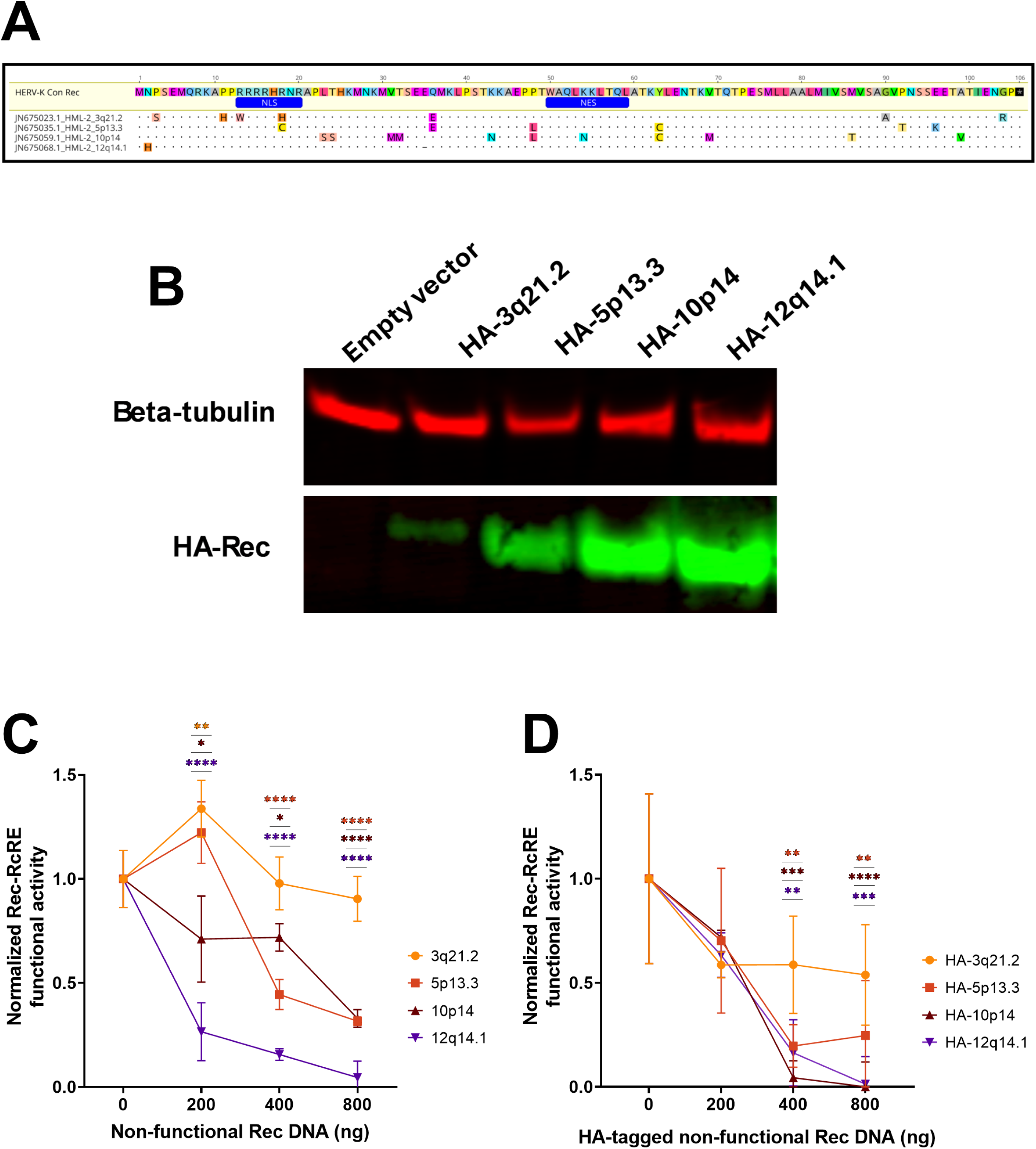
Analysis of selected non-functional HERV-K Rec proteins and their trans-dominant negative effects. (A) Multiple sequence alignment of selected non-functional Rec proteins (3q21.2, 5p13.3, 10p14, and 12q14.1) to the reference HERV-K Con Rec. Selected variants contain fewer than 15 amino acid changes compared to the consensus sequence. Amino acid differences are highlighted in color and shown in uppercase letters. Dots indicate residues identical to HERV-K Con Rec; dashes represent deletions. Position numbers correspond to HERV-K Con Rec sequence. Functional domains are annotated in blue (NLS: nuclear localization signal; NES: nuclear export signal). (B) Western blot analysis of non-functional Rec protein expression. Upper panel: β-tubulin loading control; lower panel: HA-tagged Rec protein detection. A Western blot representative of two independent experiments using two different plasmid preparations is shown. (C) Trans-dominant negative activity analysis using untagged Rec variants in 293T/17 RcRE reporter cells. Cells were co-transfected with a constant amount of HERV-K Con Rec (100 ng) and increasing amounts (0–800 ng) of non-functional Rec-expressing plasmids, maintaining total DNA at 2000 ng with empty vector. Control samples received 100 ng HERV-K Con Rec plus 1900 ng empty vector. GFP expression was measured by flow cytometry 72 hours post-transfection and normalized, with the control sample set to 1.0. Rec-RcRE functional activity was plotted as mean fluorescence intensity (MFI) of GFP. Data represent the mean value ± SD from three independent experiments. Statistical significance was assessed by ordinary Two-Way ANOVA with Dunnett’s multiple comparisons post-hoc test (*p<0.05, **p<0.01, ****p<0.0001). (D) Trans-dominant negative activity analysis using HA-tagged Rec variants in 293T/17 RcRE reporter cells. Cells were co-transfected with HERV-K Con Rec (100 ng) and increasing amounts (0–800 ng) of HA-tagged non-functional Rec variants. Data was collected, analyzed and normalized as described in (C). Data represent the mean value ± SD from three independent experiments. Statistical significance assessed by ordinary Two-Way ANOVA with Dunnett’s multiple comparisons post-hoc test (**p<0.01, ***p<0.001, ****p<0.0001).

To analyze if these Rec proteins were trans-dominant negative, we next co-transfected increasing amounts (0-800ng) of either the tagged or the non-tagged plasmids into RcRE reporter cells, with a constant amount (100 ng) of the plasmid expressing the active non-tagged HERV-K Con Rec. An appropriate amount of empty vector was added to maintain a total of 2µg of transfected DNA. We then measured Rec functional activity using flow cytometry.

In the experiment with both the non-tagged and HA-tagged Rec proteins, three of the four non-functional Rec proteins (5p13.3, 10p14, and 12q14.1) exhibited some degree of trans-dominant negative effect (Figures 4C and 4D). However, Rec 3q21.2 showed no significant inhibitory effect. Interestingly, the 12q14.1 Rec protein that displayed a very potent trans-dominant negative effect had only two changes compared to HERV-K Con Rec: an N2H mutation and a deletion at position 34 (E34del) (Figure 4A).

### Mutational Analysis of the Trans-dominant Negative Rec 12q14.1

Since Rec 12q14.1 demonstrated a potent trans-dominant negative effect while having only two amino acid changes compared to the HERV-K Con Rec, we sought to determine if one or both changes were responsible for the trans-dominance. We thus generated two variant mutants of this Rec protein as shown in Figure 5A: Variant 1, which restored a glutamic acid at position 34 (del34E), and Variant 2, which changed the histidine at position 2 to the asparagine present in HERV-K Con Rec (H2N). The variants sequences were cloned into the MSCV vector upstream of the IRES-eBFP2 region, and HA-tagged versions of the original 12q14.1 and the two variants were also generated to enable analysis of protein expression. Western blot analysis of 293T reporter cells using the HA-tagged versions of these constructs showed that all three proteins were well expressed (Figure 5B).

**Figure 5:**
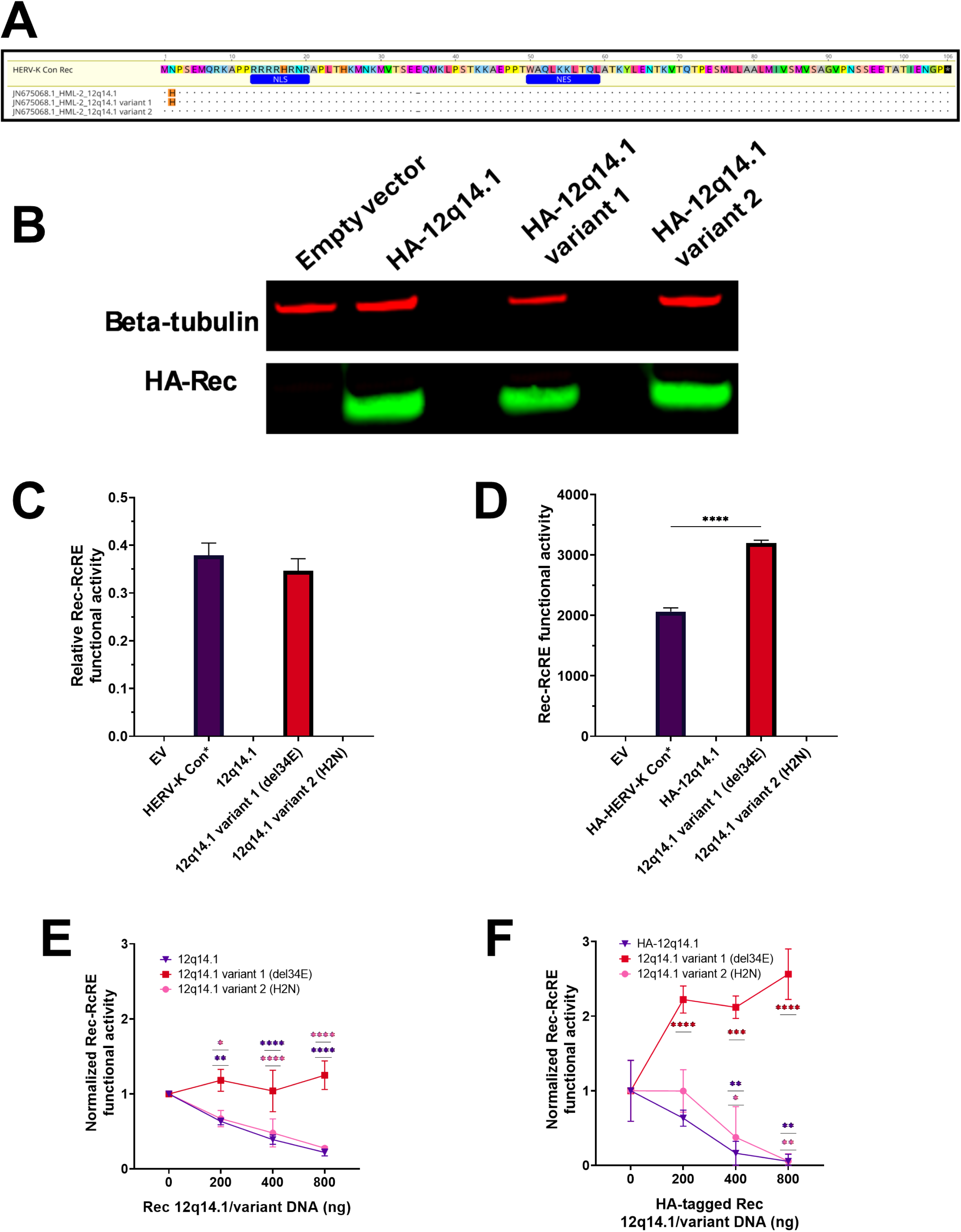
Mutational Analysis of the Trans-dominant Negative Rec 12q14.1. (A) Sequence alignments of Rec 12q14.1 variants relative to the HERV-K Con Rec sequence. The alignments show the HERV-K Con Rec sequence, the original Rec 12q14.1, and the two generated variants: Variant 1 (del34E) and Variant 2 (H2N). Amino acid differences are highlighted in color and shown in uppercase letters. Dots represent identical residues to the consensus sequence; dashes indicate deletions. Position numbers correspond to the HERV-K Con Rec consensus sequence. Functional domains are annotated in blue (NLS: nuclear localization signal; NES: nuclear export signal). (B) Western blot analysis of Rec variant protein expression. Upper panel: β-tubulin loading control; lower panel: HA-tagged Rec protein detection. A Western blot representative of two independent experiments using two different plasmid preparations is shown. (C) Functional analysis of Rec variants in 293T/17 RcRE reporter cells. Cells were transduced with retroviral vectors expressing each variant or HERV-K Con Rec (positive control). Fluorescent protein expression was then measured by flow cytometry 72 hours post-transduction. Rec-RcRE functional activity is plotted as the ratio of GFP MFI to eBFP2 MFI. Data represent the mean value ± SD from three independent experiments. Statistical significance was assessed using ordinary One-Way ANOVA with Tukey’s multiple comparisons post-hoc test. D) Functional analysis of HA-tagged Rec variants in 293T/17 2xRcRE reporter cells. Cells were transfected with retroviral vectors expressing each HA-tagged variant or HERV-K Con Rec (positive control). GFP expression was measured by flow cytometry 72 hours post-transduction. Rec-RcRE functional activity was plotted as MFI of GFP. The data represent the mean value ± SD from three independent experiments. Statistical significance assessed using ordinary One-Way ANOVA with Tukey’s multiple comparisons post-hoc test (***p<0.001). (E) Trans-dominant negative activity analysis using untagged Rec variants. 293T/17 2xRcRE reporter cells were co-transfected with HERV-K Con Rec (100 ng) and increasing amounts (0– 800 ng) of untagged variants, keeping total DNA at 2000 ng with empty vector. The data were collected, analyzed and normalized as described in Figure 4C. The data represent the mean value ± SD from three independent experiments. Statistical significance was assessed by ordinary Two-Way ANOVA with Dunnett’s multiple comparisons post-hoc test (*p<0.05, **p<0.01, ****p<0.0001). (F) Trans-dominant negative activity analysis using HA-tagged Rec variants under identical conditions as panel (E). Cells were co-transfected with HERV-K Con Rec (100 ng) and increasing amounts (0–800 ng) of HA-tagged variants. The data were collected, analyzed and normalized as described in Figure 4C. Data represent the mean value ± SD from three independent experiments. Statistical significance assessed by ordinary Two-Way ANOVA with Dunnett’s multiple comparisons post-hoc test (*p<0.05, **p<0.01, ***p<0.001, ****p<0.0001).

The untagged constructs were then packaged into retroviral particles and used to transduce the RcRE reporter cell line, in parallel with vectors that expressed the original 12q14.1 Rec and HERV-K Con Rec as a positive control. Rec activity was measured as described in Figure 2B. This experiment revealed that Variant 1 showed functional activity similar to HERV-K Con Rec (Figure 5C), whereas Variant 2 and the original 12q14.1 Rec were non-functional.

We also tested the activity of the HA-tagged 12q14.1 Rec variants. This was done by transfection of the HA-tagged plasmids into the RcRE reporter cell line and measurement of GFP MFI by flow cytometry. Similarly to the untagged constructs, HA-tagged Variant 1 showed Rec-RcRE functional activity, but the HA-tagged Variant 2 and the original HA-tagged 12q14.1 Rec did not (Figure 5D).

These results indicate that deletion of E34 abolishes Rec functional activity, whereas the H2N mutation has minimal impact.

We next analyzed the trans-dominant negative activities of the two variants by co-expressing increasing amounts of each variant with a constant amount of HERV-K Con Rec as described above for Figure 3. Variant 2 (E34del) exhibited a trans-dominant negative effect nearly identical to the original Rec 12q14.1 (Figure 5E). The HA-tagged versions gave similar results (Figure 5F). Notably, both tagged and untagged Variant 1 showed no trans-dominant negative effect, and expression of this variant actually increased the overall activity. Based on these experiments, we conclude that the glutamic acid at position 34 (E34) is essential for Rec functional activity, and that this deletion alone causes a trans-dominant negative phenotype.

## Discussion

In this study, we show that 23 of the HERV-K type 2 proviruses in the human genome have open reading frames capable of producing full-length Rec proteins. However, only 7 of these loci express functional Rec proteins in conjunction with a functional Rec-response element. Among these 7 loci, five (6q14.1, 7p22.1ab, 8p23.1a,11q22.1, and 19p12b) are polymorphic in human populations, while two (10q24.2 and 19q11) are not polymorphic at the integration site, although 19q11 has been reported as internally polymorphic (37–41). These functional loci all have nucleotide sequence changes in the *rec* gene relative to the prototypical HERV-K Con *rec* gene commonly used in published studies (38–41). However, four of these loci (6q14.1, 7p22.1ab, 11q22.1 and 19q11) express a protein with the same amino acid Rec sequence as HERV-K Con, whereas three (8p23.1a,11q22.1 and 10q24.2) display some amino acid changes (see Figure 1C). 8p23.1a only has one change (P92L) compared to HERV-K Con Rec, whereas 19p12b has two amino acid changes (A89T, E96D). None of these changes are in the known NLS and NES domains. In contrast, 10q24.2, which shows the highest Rec activity, has two changes in the NLS (R18H and N19S) and one additional change (M86T). This highlights that a few amino acid changes in retroviral regulatory proteins can significantly affect function, as shown previously for HIV Rev (42).

The evolutionary age of the HERV-K proviruses differs significantly. The relatively younger proviral insertions, such as those at loci 6q14.1 and 19p12b, are more likely to retain functional Rec proteins due to fewer accumulated mutations (1, 2). Conversely, older proviruses are typically heavily mutated, rendering their encoded Rec proteins non-functional, as seen for proviruses at several well-characterized loci (3, 5, 11). Importantly, some older proviruses, like the nearly intact loci, 10q24.2 and 19q11, maintain the potential to produce functional Rec and other viral proteins despite their age, indicating some exceptions to this (7, 11, 16).

About half of the HERV-K proviruses are type 1 and contain a deletion in Env and the overlapping Rec region. This eliminates the ability to synthesize a Rec protein, making these proviruses unable to export their unspliced genome mRNA, which requires the help of an export protein, to express Gag and GagPol. Additionally, the singly spliced mRNA, which encodes a N-terminally truncated Env protein, still retains an intron, so it is likely that Env expression would also be impaired. Thus, cells transcribing RNA from only type 1 viruses would be limited to expressing only the Np9 protein that is translated from a Rec-independent multiply spliced mRNA, unless Rec protein expression from a type 2 HERV-K encoding a functional Rec protein is expressed in the same cell. Similarly, type 2 proviruses lacking a functional Rec could also be complemented in this way to all export of their mRNAs with retained introns and expression of their viral proteins. This *trans*-complementation would be similar to our previous study, in which showed that the HIV Rev protein can complement Rec function when supplied *in trans*, to enable nuclear export of endogenously expressed HERV-K RNAs with retained introns (26).

Several studies have implicated HERV-K in various malignancies and autoimmune disorders (12, 27, 28). For instance, a recent study reported robust transcription from certain HERV-K loci in lung cancer, highlighting the potential clinical significance (15, 43). However, most of these studies have focused on expression of Env proteins (44–46). Additionally, none of these studies have examined whether Rec proteins capable of promoting nucleo-cytoplasmic RNA export were expressed.

As mentioned in the introduction, a previous study implicated HERV-K Rec proteins in cancer-related processes through interactions with the promyelocytic leukemia zinc-finger protein (PLZF), leading to c-myc upregulation and enhanced cell proliferation (12). In addition, Rec protein interactions with the human small glutamine-rich tetratricopeptide repeat-containing protein (hSGT) were reported to increase androgen receptor activity, potentially contributing to oncogenesis (28). It remains to be determined how many of the Rec proteins can interact with these proteins and cause these effects, since this may not require the protein to function in RNA export.

In addition to the few loci that can express a Rec protein that promotes RcRE-dependent RNA export, we have determined that several Rec proteins show trans-dominant negative activity. The most striking example is the Rec encoded by provirus 12q14.1, which differs from the consensus by only two amino acid changes (N2H and E34del), yet potently inhibits functional Rec activity. Our mutational analyses revealed that reintroducing glutamic acid at position 34 restored export function. Interestingly, the deletion is not in a region known to be important for nuclear export, import, or dimerization, and thus it inhibits Rec function through an unknown mechanism (15, 47). This is in contrast to the well-studied trans-dominant negative Rev protein (RevM10) that has mutations in the nuclear export signal (48–50).

The trans-dominant negative Rec proteins may serve as a natural “brake” in HERV-K replication and expression (1, 2). This could potentially limit pathogenic HERV-K effects, and these Rec variants may thus have been positively selected during HERV-K evolution and spread. In addition, the deletion of Rec in the type 1 HERV-K proviruses eliminates potential pathogenic effects due to Rec overexpression. However, since Rec can act in *trans*, if expressed from a type 2 virus that is active in the same cell, type 1 viruses do not need to encode Rec for replication.

Our discovery that active proviral copies of HERV-K can encode both active and inhibitory Rec proteins shows the complexity of Rec regulation. Levels of HERV-K expression and potential host cell effects of Rec activity will vary depending on which HERV-K loci are active, and it remains possible that Rec is an important factor in some human cancers. Further studies of the oncogenic properties of Rec and how this relates to the function of Rec as a nuclear export factor will be needed to elucidate this.

## Materials and methods

### Identification of HERV-K HML-2 Rec ORFs

Human endogenous retrovirus K (HERV-K) HML-2 proviral sequences (n=91) were retrieved from the NCBI Data Repository (GenBank ID JN675007-JN675097) (5). Each HERV-K HML-2 genome was aligned to the annotated HERV-K Con reference genome using Geneious Prime software (version 2020.2, Biomatters Ltd., Auckland, New Zealand) to transfer annotated proviral features. Type 1 proviruses were identified based on the presence of the 292bp deletion in the pol-env region and the GA-GT mutation in the 5’ splice site. Rec coding sequences from the identified type 2 proviruses lacking the 292bp deletion were generated by extracting and joining the annotated Rec exons from each type 2 provirus. Sequence alignments were performed using the MUSCLE algorithm with default parameters as implemented in Geneious.

### Plasmids and Vector Construction

#### Reporter Vector System

The dual-color Rec-RcRE reporter vector was adapted from the previously described Rev-RRE HIV reporter system (35, 36). The HIV RRE sequence was replaced with two tandem copies of the HERV-K RcRE (2xRcRE) while maintaining the same vector backbone and fluorescent protein reporters. The RcRE sequence was derived from the (GenBank AF179225.1) and synthesized by Integrated DNA Technologies (IDT, Coralville, IA, USA).

#### Retroviral Vectors

For Rec protein expression, HERV-K Rec open reading frames (ORFs) were commercially synthesized (Genscript, Piscataway, NJ, USA) and cloned into murine stem cell virus (MSCV)-based retroviral vectors containing an IRES-eBFP2 cassette (pMSCV-IRES-eBFP2). Cloning was performed using standard restriction enzyme digestion (EcoRI, XhoI; New England Biolabs, Ipswich, MA, USA) followed by T4 DNA ligase-mediated ligation (Thermo Fisher Scientific, Waltham, MA, USA).

To generate N-terminal HA-tagged Rec expression constructs, we PCR-amplified each Rec variant using a forward primer encoding a 5′ HA epitope tag (MYPYDVPDYA) and a Kozak consensus sequence immediately upstream of the Rec start codon. This primer also contained a 17 bp overhang complementary to the MSCV vector backbone. The reverse primer annealed near the 3′ end of the Rec open reading frame and included a complementary overhang to the MSCV vector. PCR was performed using a high-fidelity DNA polymerase (Thermo Fisher Scientific) with the following cycling conditions:

Initial denaturation: 94°C 30 seconds

25 cycles of: 94°C for 15 seconds, 65°C for 30 seconds, 68°C for 30 seconds

Final extension: 68°C for 10 minutes

The resulting PCR products were cloned into a modified MSCV vector lacking the IRES-eBFP2 cassette using Gibson Assembly (NEB). All constructs were verified by Sanger sequencing. The primer sequences are listed in Supplemental Table S2.

#### Mutational Analysis Constructs

Rec 12q14.1 variants were generated using synthetic double-stranded DNA fragments (gBlocks; Integrated DNA Technologies) and subsequently cloned into the pMSCV vector upstream of the IRES-eBFP2 cassette via Gibson assembly following the manufacturer’s protocol. Each plasmid was assigned a unique identifier (pHRXXXX) as detailed in Supplementary Table S1. Construct integrity was confirmed by Sanger sequencing (Eton Bioscience Inc).

### Cell Culture and Viral Vector Production

#### Cell Maintenance

293T/17 cells were maintained in Iscove’s Modified Dulbecco’s Medium (IMDM; Gibco, Thermo Fisher Scientific) supplemented with 10% bovine calf serum (BCS, Life Technologies) and 50µg/ml gentamicin (Gibco). Cells were cultured at 37°C in a humidified 5% CO_2_ incubator.

#### Retroviral Vector Production

Retroviral vectors expressing Rec variants were produced in 293T/17 cells. Briefly, 8×10^6^ cells were seeded in 15cm plates containing 20ml growth medium (IMDM, 10% BCS, 50µg/ml gentamicin). After 24 hours, cells were transfected in serum-free conditions using polyethyleneimine (PEI; Polysciences, Warrington, PA, USA 1 mg/ml stock solution, pH 7.0) with a plasmid mixture containing 30µg pMSCV-Rec construct, 12.84µg MLV Gag-Pol (pHIT) plasmid, and 5.16µg VSV-G expression plasmid (pMD2.G, Addgene). For transfection, DNA was mixed with PEI at a ratio of 1:3 (DNA:PEI) in serum-free IMDM and incubated for 20 minutes at room temperature before adding to cells. Following a 6-hour incubation in serum-and antibiotic-free IMDM, the transfection medium was replaced with fresh growth medium.

Forty-eight hours post-transfection, the viral supernatant was collected and cleared by centrifugation (380 RCF, 4°C, 5 minutes) to remove cellular debris. The cleared supernatant was then aliquoted (1 ml per aliquot) and stored at -80°C until use. Lentiviral particles containing the tandem RcRE reporter were generated similarly, except using psPAX2 (Addgene) as the packaging plasmid in a ratio of 12.84µg psPAX2, 30µg reporter plasmid, and 5.16µg pMD2.G per 15cm plate.

For viral titration, serial 10-fold dilutions were prepared in serum- and antibiotic-free culture medium. 293T/17 cells (3×10^4^) were seeded in 96-well plates 24 hours before transduction. Cells were transduced with 100µl of diluted viral stocks containing 6µg/ml DEAE-dextran (Sigma-Aldrich) and incubated for 6 hours at 37°C, 5% CO2. Following medium replacement, cells were cultured for 72 hours, then harvested and resuspended in PBS containing 5% BCS. mCherry or eBFP2 expression was quantified using an Attune NxT flow cytometer with autosampler (Thermo Fisher Scientific). Viral titers were calculated as transducing units per ml (TU/ml) based on the percentage of fluorescent protein-positive cells in the linear range of dilutions (between 1% and 20% positive cells).

### Generation of Stable Reporter Cell Line

A stable 293T/17 reporter cell line (293T/17: pNL4-3(GFP)(HERVK2xRcRE) (mCherry)) was established through lentiviral transduction. Briefly, 3×10^6^ 293T/17 cells were seeded in 10cm dishes 24 hours before transduction. Cells were transduced at MOI 0.1 with the lentiviral reporter construct in serum- and antibiotic-free IMDM containing 6µg/ml DEAE-dextran.

Following 6-hour incubation at 37°C, 5% CO2, the virus-containing medium was replaced with standard growth medium.

Transduced cells were subjected to fluorescence-activated cell sorting (FACS) using a BD Influx System (BD Biosciences, San Jose, CA, USA). Single cells expressing the highest levels of mCherry were isolated and seeded into individual wells of 96-well plates containing IMDM supplemented with 20% BCS and 50 µg/ml gentamicin. After a two-week expansion period, clonal populations were cryopreserved in growth medium containing 10% DMSO (Sigma-Aldrich). The established clonal cell line was subsequently evaluated for consistent and robust reporter gene expression. Functional assays were performed to confirm the responsiveness of the reporter system to the presence of functional Rec proteins, indicated by GFP expression upon Rec-RcRE interaction.

### Functional Analysis of Rec Variants

To assess Rec-RcRE functional activity, reporter 293T/17 cells (3×10^4^ cells/well) were seeded in 96-well plates 24 hours before transduction. Cells were transduced at MOI 1 with retroviral vectors expressing individual Rec variants in serum- and antibiotic-free RPMI containing 6µg/ml DEAE-dextran. Following 6-hour incubation at 37°C, 5% CO2, virus-containing medium was replaced with standard growth medium. At 72 hours post-transduction, cells were harvested by trypsinization using 0.25% Trypsin-EDTA (Gibco), resuspended in PBS containing 5% BCS, and analyzed by flow cytometry. Single-color controls were included for proper compensation while measuring mCherry, GFP, and eBFP2 expression.

### Western Blot Analysis

To analyze protein expression levels, reporter 293T/17 cells were transfected with 2µg of MSCV plasmids expressing either functional or non-functional Rec variants using Lipofectamine 3000 (Invitrogen, Thermo Fisher Scientific) according to the manufacturer’s protocol. At 72 hours post-transfection, cells were harvested and split for parallel flow cytometry analysis and Western blot.

For Western blot samples, cells were washed three times with cold PBS and lysed in buffer containing 50mM Tris pH 7.4, 150mM NaCl, 1.5% SDS, and protease inhibitors. Lysates were passed through a 21G needle (BD), heat-treated (90°C, 10 min), and cleared by centrifugation (14,000 RCF, 10 min) using a microcentrifuge (Eppendorf, Hamburg, Germany). Samples were prepared for SDS-PAGE by combining 50µl lysate with NuPAGE LDS Sample buffer (4X) (Invitrogen) and NuPAGE Sample Reducing Agent (10X) (Invitrogen), followed by heating at 70°C for 10 min.

Proteins were separated on Novex 4-12% Bis-Tris gels (Invitrogen) at 120V for 1.5 hours using Electrophoresis System and transferred to PVDF membranes at 30V for 1 hour at 4°C).

Membranes were blocked with 5% BSA (Sigma-Aldrich) in TBS (1 hour, room temperature) and incubated overnight at 4°C with primary antibodies (Mouse anti-HA Tag, Cell Signaling Technology; 1:30000 dilution) and (Rabbit anti-Beta Tubulin Cell Signaling Technology; 1:2000 dilution) diluted in TBS-T (TBS + 0.1% Tween 20) containing 5% BSA.

Membranes were washed three times with TBS-T (10 minutes each) and were incubated with IRDye-conjugated secondary antibodies (donkey anti-mouse IRDye 800CW, LI-COR Biosciences; 1:30000 dilution) and (goat anti-rabbit IRDye 680RD; LI-COR Biosciences, 1:5000) for 1 hour at room temperature. After three additional washes with TBS-T, protein detection was performed using a LICOR Odyssey CLx scanner, and band intensities were quantified using Image Studio Software (version 5.2, LI-COR Biosciences). For quantification, Rec protein band intensities were normalized to beta-tubulin loading controls from the same lanes. Antibodies used are listed in Supplementary Table S3.

To normalize functional activity to protein expression, we divided the average mean fluorescence intensity (MFI), measured by flow cytometry, by the relative protein abundance of each Rec variant as detected by Western blotting of the HA-tagged protein. To accurately compare protein levels between samples, HA-tag signals were first normalized to β-tubulin as a loading control. The resulting normalized activity was calculated as follows:

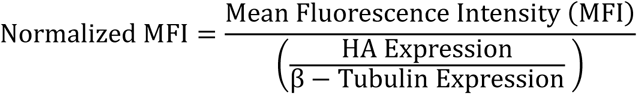

Where:

- **MFI** = Average Mean Fluorescence Intensity measured by flow cytometry, representing functional activity
- **HA** = Average HA-tagged Rec protein level measured by Western blot
- **β-Tubulin** = Loading control used to normalize HA signal across samples

### Flow cytometry

Cells were harvested by trypsinization and resuspended in PBS containing 5% BCS. Samples were analyzed using an Attune NxT flow cytometer with autosampler (Thermo Fisher Scientific). For each sample, a minimum of 30,000 events were collected. Single-color controls were used for compensation using the automated compensation feature of the FlowJo software. Data analysis was performed using FlowJo v10.6.1 (FlowJo, LLC, BD Biosciences).

The analysis workflow included initial gating on single cells using forward and side scatter properties (FSC-A vs. SSC-A) followed by doublet exclusion (FSC-A vs. FSC-H). Gates for mCherry, eBFP2, and GFP-positive populations were established using untransduced 293T/17 cells as negative controls. All subsequent analyses were performed on the mCherry-positive population.

Relative Rec-RcRE functional activity was quantified as follows:

I. For Figures 2B and 5C. Calculated as the ratio of GFP to eBFP2 mean fluorescence intensity (MFI)
II. For all other figures: Measured as GFP MFI from mCherry-positive populations

### Trans-dominant negative Rec assays

To measure the effect of co-expressing the functional HERV-K Con with the selected non-functional Rec proteins, 2.5 x 10^5^ reporter 293T/17 cells were seeded in 24-well plates and transfected with 100ng of the pMSCV-HERV-K Con Rec, and either 0, 200, 400 or 800ng of pMSCV-IRES-eBFP2 vector expressing each of the four test non-functional Recs. Transfections were performed using Lipofectamine™ 3000 Transfection Reagent (Invitrogen) according to the manufacturer’s protocol. For each well, DNA was mixed with 1μl of the P3000 and 1.5 μl of Lipofectamine 3000 in 50 μl of Opti-MEM. Variable amounts of an empty pMSCV plasmid were added to maintain a constant DNA mass of 2 μg in each transfection. Cells were incubated for 72 hours and subjected to flow cytometry to measure the expression of mCherry and GFP. The same method was used to test the effects of co-expressing HERV-K Con Rec with the Rec 12q14.1 point mutants.

### p24 assays

To assess Rec-mediated viral protein expression, 2.3×10^5^ 293T/17 cells were seeded per well in 12-well plates. Cells were transfected with a constant amount (1500 ng) of Gag-Pol RcRE reporter vector and increasing amounts of Rec expression plasmids (50, 100, 150, and 250 ng). Transfections were performed using PEI as described above. At 72 hours post-transfection, culture supernatants were harvested, centrifuged at 300 × g for 5 minutes to remove cellular debris and analyzed for p24 capsid protein using an in-house ELISA assay as previously described (51).

Briefly, 96-well plates (Nunc MaxiSorp, Thermo Fisher Scientific,) were coated with 100 μl of anti-p24 monoclonal antibody (NIH AIDS Reagent Program) at 1:10 000 dilution in PBS overnight at 37°C. Plates were washed five times with PBS and blocked with 200 μl of PBS containing 5% Blocking buffer (PBS with 5%BCS) for 1 hour at 37°C. After washing, 100 μl of culture supernatants or p24 standards (NIH AIDS Reagent Program) ranging from 12.5 to 1600 pg/ml were added to the wells and incubated for 2 hours at 37°C. Plates were washed and incubated with 100 μl of mouse anti-p24 polyclonal antibody (NIH AIDS Reagent Program) at 1:10000 dilution for 1 hour at 37°C. After washing, 100 μl of horseradish peroxidase-conjugated goat anti-mouse IgG (Abcam) at 1:20 000 dilution was added and incubated for 30 min at 37°C. Plates were washed and developed with 100 μl of substrate for 30 minutes at room temperature in the dark. The reaction was stopped with 50 μl of 1N H_2_SO_4_, and absorbance was measured at 450 nm using a microplate reader (BioTek Synergy HTX, BioTek Instruments). A four-parameter logistic regression standard curve was generated and p24 concentrations in the samples were interpolated from this standard curve. Each sample was assayed in duplicate, and the average value was reported.

### Statistical analysis

Statistical Analysis: Statistical analyses were performed using GraphPad Prism software (version 9.0; GraphPad Software, San Diego, CA, USA). Data are presented as mean ± standard deviation (SD), derived from at least three independent experiments unless otherwise indicated in the figure legends. For comparisons involving more than two experimental groups, statistical significance was determined using ordinary one-way ANOVA followed by Tukey’s multiple comparisons test, or by non-parametric one-way ANOVA with Dunn’s multiple comparisons post-hoc test, depending on the outcome of data normality assessments. For comparisons between two groups, either an unpaired two-tailed Student’s t-test or the non-parametric Mann–Whitney U test was used, as appropriate. Trans-dominant negative activity assays, involving multiple conditions across different expression levels, were analyzed using ordinary two-way ANOVA followed by Dunnett’s multiple comparisons test against a single control (functional Rec alone), assuming pooled variance. For experiments involving multiple comparisons, p-values were adjusted using the Benjamini-Hochberg procedure to control the false discovery rate (FDR). A p-value of less than 0.05 was considered statistically significant. Significance thresholds used throughout the manuscript were defined as follows: *p<0.05, **p<0.01, ***p<0.001, ****p<0.0001. The specific statistical tests applied to individual experiments are indicated in the corresponding figure legends. Graphs were prepared using GraphPad Prism.

### c

The monoclonal anti-HIV-1 p24 antibody (183-H12-5C, ARP-3537) used in the p24 ELISA was obtained from the NIH HIV Reagent Program, Division of AIDS, NIAID, and was contributed by Dr. Bruce Chesebro and Kathy Wehrly. The flow cytometry data for this manuscript were generated at the University of Virginia Flow Cytometry Core Facility (RRid:SCR_017829), with partial support from the NCI Cancer Center Grant (P30-CA044579). Partial salary support was provided by the Myles H. Thaler Endowed Professorship (D.R.), and the Charles Ross Jr Endowed Professorship (M.-L. H) at the University of Virginia. Partial stipend support was provided by NIH National Cancer Institute Center Grant P30-CA044579 (K.Z). Research support was provided by Myles H. Thaler Research Support Gift to UVA and grant R01-CA206275 from the National Cancer Institute (NIH).

